# De novo design of a macrocycle induced dimerization system for cellular control

**DOI:** 10.64898/2026.04.24.720480

**Authors:** Stephanie Hanna, Patrick J. Salveson, Basile Wicky, Madison A. Kennedy, Derrick R Hicks, Carolina Moller, Suna Cheng, Xinting Li, Mohamad Abedi, Brian Coventry, Meerit Y. Said, Asim K. Bera, Alex Kang, Barry L. Stoddard, David Baker

## Abstract

Investigating and manipulating cellular events requires precise control of protein function. To enable control over cellular processes, we set out to design a chemically induced dimerization (CID) system consisting of a *de novo* designed ligand and protein pair. Here we describe the design of a C2 symmetric membrane permeable macrocyclic peptide and a cognate protein homodimer which binds the macrocycle through a large interface with both chains. The designed homodimer binds the macrocycle with a *K*_D_ of 36 nM, and the x-ray crystal structure of the protein homodimer-macrocycle complex is very close to the computational design model, with the C2 axis of the macrocycle aligned with the homodimer C2 axis. Transcriptional and split luciferase assays in mammalian cells demonstrates conditional control over both a reporter gene expression and luciferase reconstitution.

## Introduction

Chemically induced dimerization (CID) systems in which protein-protein association is dependent on membrane permeable small molecule ligands are broadly useful in synthetic biology to enable external control of cellular processes. CID systems adapted from nature have been used to control biological processes ranging from apoptosis to targeted protein degradation to transcription^1,2,3^. The power and limitations of such systems are exemplified by the rapamycin-FKBP-FRB system which is widely used for cellular control. Rapamycin not only effectively induces FKBP-FRB association but also is an immunosuppressant that dimerizes endogenous proteins, leading in some cases to undesired off-target effects *in vivo*^4^. Alleviation of the toxicity issues associated with rapamycin CID systems have been accomplished either by derivatization of rapamycin and FK506 and the FKBP protein, or adapting CID systems native to plants into mammalian cells^5, 6, 7, 8^. Expansion to other ligand classes has relied upon directed evolution approaches to engineer a binding interface for a desired ligand or finding antibody fragment binding partners through phage display for a ligand or existing protein ligand binary complex^9, 10, 11, 12, 13^. Protein design methods can in principle yield a variety of CID systems; a farnesyl pyrophosphate sensor relying on a maltose binding and an ankyrin repeat protein, as well as a CID composed of NS3a protease, grazoprevir, danoprevir, and asunaprevir along with three designed proteins are two such examples^14, 15^. Developing CIDs with new chemical ligands, however, has been relatively unexplored due to a lack of protein domains that bind disparate or xenobiotic molecules. An attractive class of possible compounds for CID system construction are sets of designed, structured macrocycles (MCs) that are chemically diverse, easy to synthesize, and membrane permeable. These compounds have considerable exposed surface area available for binding, and an almost unlimited number of such molecules can now be designed^16^.

We reasoned that designed proteins that dimerize in the presence of MCs could yield a wide variety of CID systems. To highlight the biochemical diversity that might be possible, we designed both the MCs and the sensor protein from scratch.

## Results

For the MC component of the system, desirable properties are 1) membrane permeability for intracellular applications, 2) sufficient exposed surface area to drive protein association, and 3) a well-ordered tertiary structure to favor on-target while disfavoring off-target interactions. Guided by previous studies indicating that structured MCs are generally cell permeable if all polar groups are satisfied in hydrogen bonding interactions, we develop a method to generate closed MCs with internal backbone-hydrogen bonding^16^. We generate large sets of loops of 5, 6, and 7 residues in which the terminal residues make beta-strand like hydrogen bonds and the remaining residues were N-methylated. Closed MCs are constructed by identifying pairs of loops with the terminal hydrogen bonding residues in nearly identical orientations. For computational efficiency, this identification is done using geometric hashing: for each loop we calculate the rigid body transform between the N- and C-terminal residues in both the N-to-C and C-to-N orientations, and closed MC backbones are then constructed by identifying pairs of loops for which the N-to-C transform of one is the same as the C-to-N transform of the other (Fig. 1A)^17^. These resulting MCs have two backbone-hydrogen bonds between the two residues at the fusion junctions, and the remainder of the backbone amides were N-methylated, leaving no unsatisfied backbone amides to compromise membrane permeability. For each such generated structure, we use RosettaFastDesign to generate low-energy amino acid sequences allowing both L and D amino acids at each position while maintaining the pattern of N-alkylation to maintain full NH satisfaction throughout the MC (Fig. 1B).

**Figure 1.**
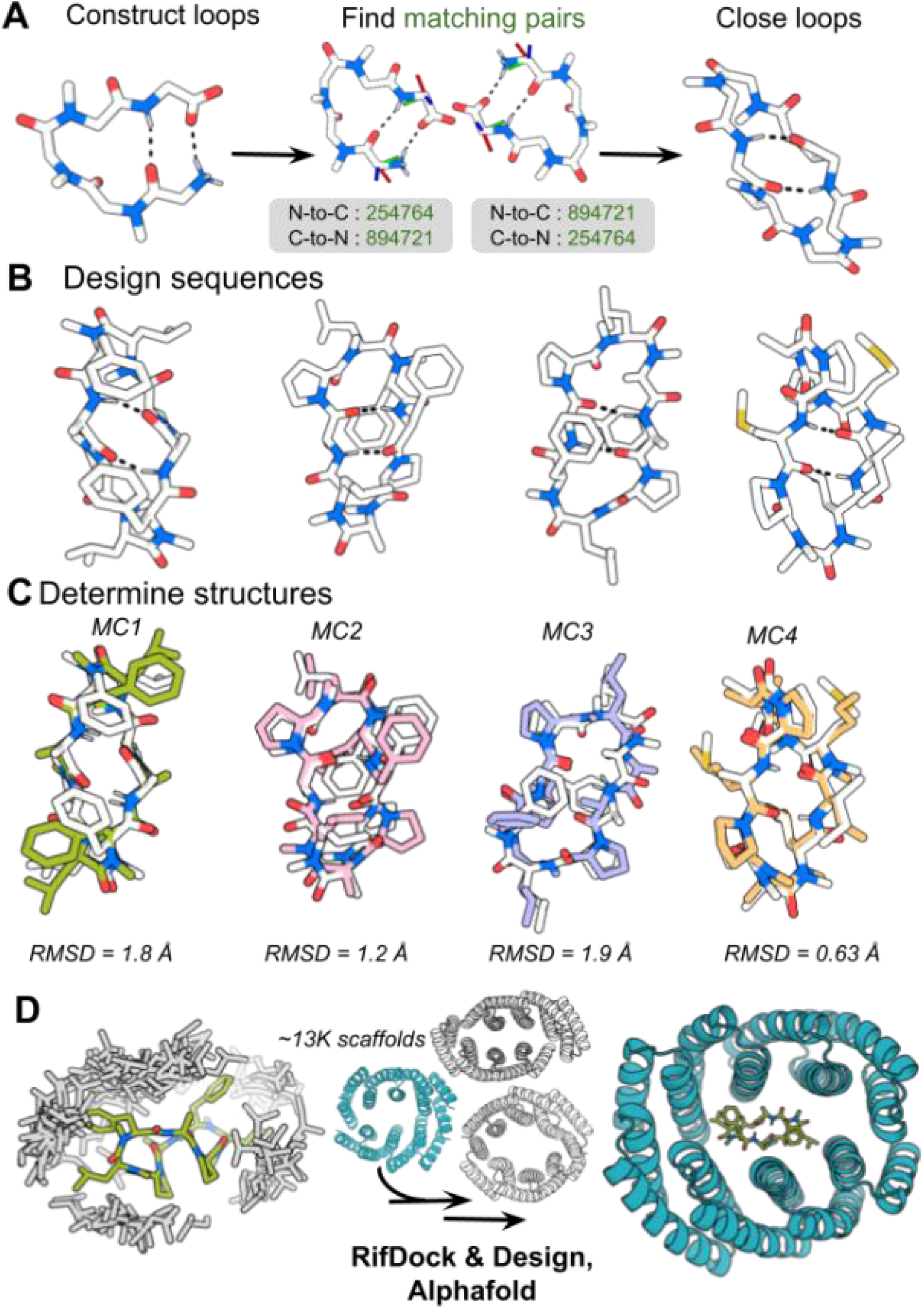
Computational design procedure. **a** Five, six, or seven residue loops were constructed to mimic beta hairpins. Construction of macrocyclic peptide backbones is shown using a representative, five residue starting loop. 6D transforms were calculated and binned. Pairs of loops were fused together if their termini geometry match, as indicated by green bin numbers. **b** RosettaFastDesign was used to generate sequences for macrocyclic peptides, allowing both L and D hydrophobic residues. Design models of the MCs are shown in white with dashed black lines indicating hydrogen bonds. **c** X-ray crystal structures (color) overlaid on the corresponding design models (white) for four designs. The all atom RMSDs between the design models and crystal structures are depicted beneath the MCs. **d** A rotamer interaction field was generated from the MC1 crystal structure (green); leucine side chains in the RIF process are shown for illustration (gray). RifDock was then used to place the peptide into the central cavity of homodimeric scaffold proteins. Protein peptide interactions were optimized with Rosetta fastDesign, retaining the protein-protein interfaces, and the designs were selected for experimental characterization based on AF2 and Rosetta metrics.

MC1, MC2, MC3 and MC4 are excellent candidates for induced dimerization systems as they are likely membrane permeable, with all backbone amides either engaged in hydrogen bonds or N-methylated and have well defined structures. An ordered structure reduces the chance of unwanted interactions with natural proteins and should increase affinity for designed binders since backbone entropy would not be lost in binding. The first three are asymmetric, and hence well suited to the design of induced heterodimerization CIDs. While we could apply these asymmetric macrocycles to the design of heterodimeric CIDs, this paper focuses on building induced homodimerization systems around the internally symmetric MC1 for two reasons: first many important processes in biology, such as caspase activation, involve homodimerization, and second, the design problem is simpler as only one protein chain need be designed^19^.

We solved crystal structures of four of these designed MCs, MC1, MC2, MC3, and MC4 (Fig. 1B). Three are asymmetric–MC2, MC3, and MC4– and have sequences apL*F*FPa*l*, fpL*a*APl*f*, and XPa*l*iX*Pa*P, respectively, where the asterisk represents N-methylation, X represents norleucine, capital letters are L amino acids and lower case D amino acids. As is evident in the structures, the two loops flanking the central backbone hydrogen bond pair have different conformations in different designs, and the structures are stabilized by torsional restraints from multiple proline residues as well as aromatic packing interactions (Fig. 1B). In proteins, backbone hydrogen bonds do not generally favor specific conformations as there are large numbers of alternative conformations in which these can be made, but in our designed MCs, extensive N-methylation and backbone cyclization strongly favors antiparallel versus parallel pairings. Thus, hydrogen bonding can effectively only occur between the intended residues and in the designed arrangement, which likely also contributes to structural specification. The crystal structures of these designs are very close to the design models over the peptide backbone, but there are several evident flips of aromatic residues in some cases (Fig. 1C). The design scheme can be readily extended to incorporate loops which bind protein targets of interest as a potential therapeutic strategy. The sequence of MC1, apL*F*apL*F*, is internally duplicated, and both the design model and the crystal structure have C2 symmetry. Two of the eight amides are in the cis-conformation, which has been associated with increased membrane permeability^18^. An additional three crystal structures were solved for MC5 and MC6 as well as an L4F mutant of MC6 (Supplementary Note 1). MC5 and MC6 loops were not in agreement between the design and crystal structure, but the MC6-L4F mutant and its crystal structure align closely (Supplementary Fig. 1).

For the protein component of the new system, we chose to start from designed helical repeat homodimers containing central binding pockets with a wide range of shapes and sizes, including many that are in the size range of MCs^20^. An initial round of binder design with these scaffolds and MC1 yielded a co-crystal structure where MC1 was not in the correct orientation of the binding pocket with respect to the design model (Supplementary Fig. 2). The sequences of the original scaffolds were redesigned with ProteinMPNN to increase folding robustness and used for a successive round of binder design with MC1^21^. Homodimeric binders for MC1 were computationally designed by first generating a rotamer interaction field around MC1 using RIFgen. RIFgen generates large numbers of disembodied sidechains making favorable interactions with the proposed ligand^22^. RIFdock was then used to place MC1 into the central cavity of each homodimeric scaffold to maximize the number of these favorable sidechain interactions that could be realized, resulting in RIF sidechains with N-Ca -C backbone intersecting with backbone atoms of the scaffold (Fig. 1C). Docks were filtered on interface shape complementarity, binding energy (ΔΔG), contact molecular surface (CMS) area between the protein and the peptide, and contact patch. The peptide protein interface was then redesigned with Rosetta FastDesign, leaving the sequence of MC1 fixed. The resulting designs were filtered for the number of hydrogen bonds made to the ligand, CMS, ΔΔG, interface shape complementarity, and the number of buried unsaturated hydrogen bond donors. AlphaFold 2 (AF2) structure predictions were generated for each and predictions were then filtered on predicted local distance difference test (pLDDT), and predicted template modeling score (pTM), yielding 263 designs, seventeen of which were selected for experimental validation.

Synthetic genes encoding the designed proteins were obtained, expressed in *E. coli*, and evaluated for binding to MC1 by equilibrium dialysis. An initial binding screen by equilibrium dialysis suggested one out of eleven screened designs, CID7, bound MC1 at both protein concentration points assessed (Supplementary Fig. 3). CID7 exhibited a monodisperse size exclusion chromatography (SEC) trace. We then determined a 36 nM *K*_*D*_ of homodimeric CID7 to MC1 by Isothermal titration calorimetry (ITC) (Fig. 2A-B). Mass photometry experiments with maltose binding protein fused to CID7 revealed the approximate binding affinity of the CID7 homodimer is 0.4 nM (Supplementary Fig. 4).

**Figure 2.**
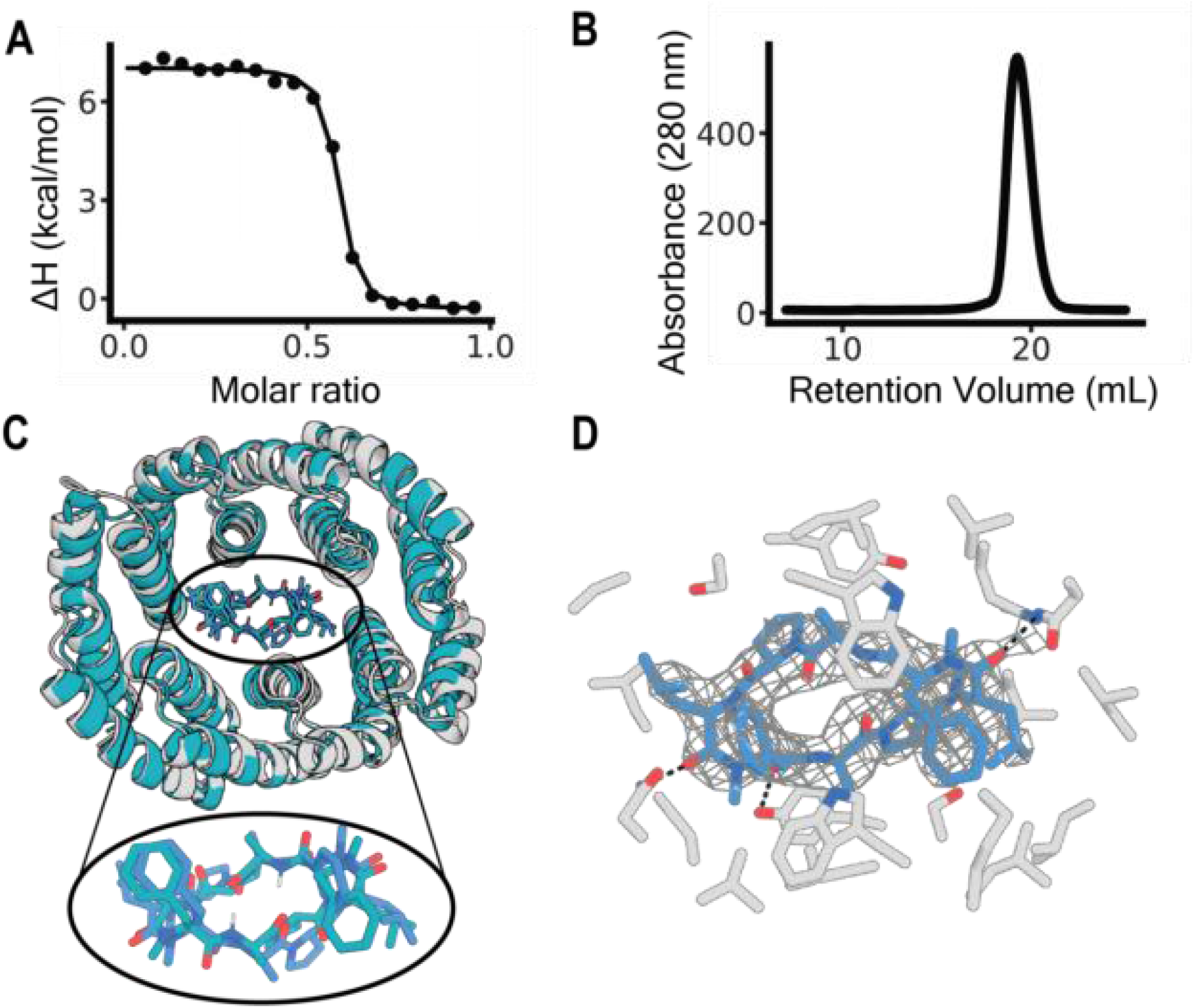
Structural and biophysical characterization of the designed macrocycle-protein interaction. **a** Titration curve starting with 250 μM of MC1 with CID7 obtained from ITC. *K*_D_ was measured at 36 nM. Data for one titration is shown. **b** CID7 SEC trace. Absorbance was measured at 280 nm. **c** Overlay of the design model (teal) with the crystal structure (gray). RMSD between the protein chains is 0.95 **Å**. Zoom of the binding pocket shows MC1 of the design model (teal) and MC1 of the crystal structure (blue). RMSD between the two structures is 0.97 **Å. d** Zoom in of the crystal structure binding pocket. Protein residues (gray) within 4 **Å** of MC1 (blue) are shown and hydrogen bonds are indicated with dashed black lines. Electron density around MC1 is shown in gray mesh with contour level 2.

The structure of MC1 in complex with CID7 was solved by X-ray crystallography. The complex has designed C2 symmetry, with the orientation of the peptide in the binding pocket very close to the design model and a 0.97 **Å** C-alpha RMSD between MC1 in the design and crystal structure (Fig. 2C). For the protein chains, the C-alpha RMSD between the design model and crystal structure was 0.95 **Å**. The proline rings of MC1 pucker in the crystal structure, allowing for a backbone carbonyl to engage in hydrogen bonding with Y89. N53 and N218 also form hydrogen bonds with MC1 backbone carbonyls (Fig. 2D).

We investigated the membrane permeability of MC1 in an artificial membrane permeability assay (PAMPA). MC1 has an apparent permeability of 1.1×10^-5^ cm/s, above the 1.5×10^−6^ cm/s cutoff for a compound to be considered permeable^16^ (Supplementary Fig. 5). We also assessed the cytotoxicity of MC1 in HEK293T cells using an LDH-Glo assay (Promega J2380). HEK293T cells treated with MC1 titrated 1 in 2 from 50 uM to 50 nM and aliquots of cell media were analyzed for LDH presence 48 h after treatment. We compared the LDH release to a vehicle only control (cell media with 1% DMSO) as well as a maximum LDH control where cells were treated with TritonX 15 minutes before LDH measurements were taken. We observed no significant cytotoxicity. Treatment with MC1 showed similar luminescence and percent cytotoxicity to that of vehicle-treated cells (Supplementary Fig. 6A-B).

We evaluated the inducibility and reversibility of the CID7 system in an in-cell split luciferase (Nanobit) assay. CID7 proteins were expressed in HEK293T cells as N-terminal fusions to either smBiT (smBiT-CID) or lgBiT (lgBiT-CID7) NanoLuc® luciferase. The luminescence of samples expressing CID7 fusions was somewhat higher than that of a negative control consisting of Halo protein fused to smBit on the C-terminus and lgBiT-CID7, suggesting some background dimerization. Upon addition of MC1, approximately 3-fold increase in luminescence was observed. In contrast, the luminescence signal decreased in cells treated with vehicle rather than MC1 (Fig 3B). After washing out the ligand and adding additional substrate, the luminescent signal returned to close to the level before treatment with MC1 or vehicle, indicating reversibility (Fig 3B).

**Figure 3.**
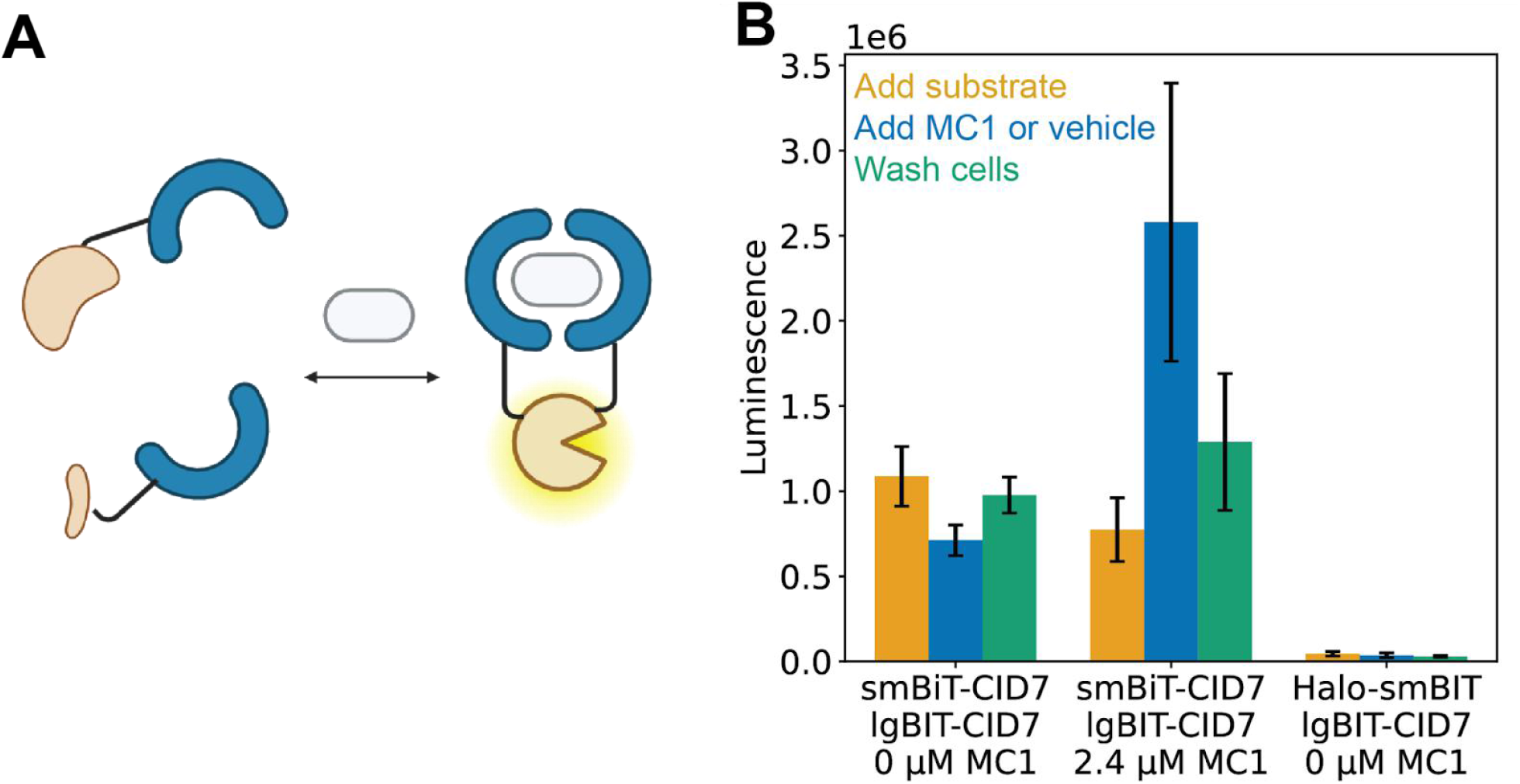
Inducibility and reversibility of MC1-dependent dimerization using split luciferase assay in HEK293T cells. **a** Schematic representation of the split luciferase assay. CID7 proteins (blue) were expressed in HEK293T cells as fusions with either small or large bit luciferase. Dimerization of the CID7 proteins reconstitutes the luciferase, which then can convert substrate to product, producing a luminescent signal. Created in BioRender. **b** HEK293T cells were transiently transfected with either LgBiT-CID7 and smBiT-CID7 or smBiT-CID7 and Halo-LgBiT. Substrate was added to cells and yellow bars represent the mean luminescence of each sample after substrate addition. Cells were then treated with either vehicle (0 μM, Opti-MEM with 2.5% DMSO) or 2.4 μM MC1, and mean luminescence was plotted (blue). Cells were washed with vehicle, additional substrate was added, and the mean luminescence was plotted (green). n=3 biological replicates from individual transfections were used for all conditions tested. Error bars represent standard deviation of the mean.

We next evaluated whether MC1 could induce dimerization of CID7 in a mammalian two-hybrid assay. CID7 was independently fused to either a zinc finger DNA binding domain (DBD) or a p65 activation domain (AD) and the two constructs were expressed in HEK293T cells along with an enhanced green fluorescent protein (eGFP) reporter which should be activated by reconstitution of the split transcription factor^23^. Upon addition of MC1, there was an increase in eGFP expression over control cells treated with cell media with 1% DMSO, likely reflecting dimerization of CID7 and reconstitution of the split transcription factor. Titration of MC1 yields a dose response curve with an EC_50_ of 9.4 μM and a 16-fold increase in geometric mean fluorescence intesity (gMFI) of eGFP at saturating ligand concentrations (Fig. 3A-B). Transfection efficiency was assessed by mscarlett-1 and mtagBFP2 fluorescence levels which are co-expressed with each construct through an internal ribosome entry site (IRES) in the AD and DBD plasmids, respectively. In the absence of MC1, there is some activation of eGFP expression due to basal homodimerization of CID7 in cells with high transfection efficiency (Supplementary Fig. 7). In control experiments, we generated similar constructs utilizing the FRB-FKBP rapamycin inducible system and observed 17-fold increase in eGFP gMFI between control cells and the highest concentration of rapamycin, 10 μM (Fig. 4C-D).

**Figure 4.**
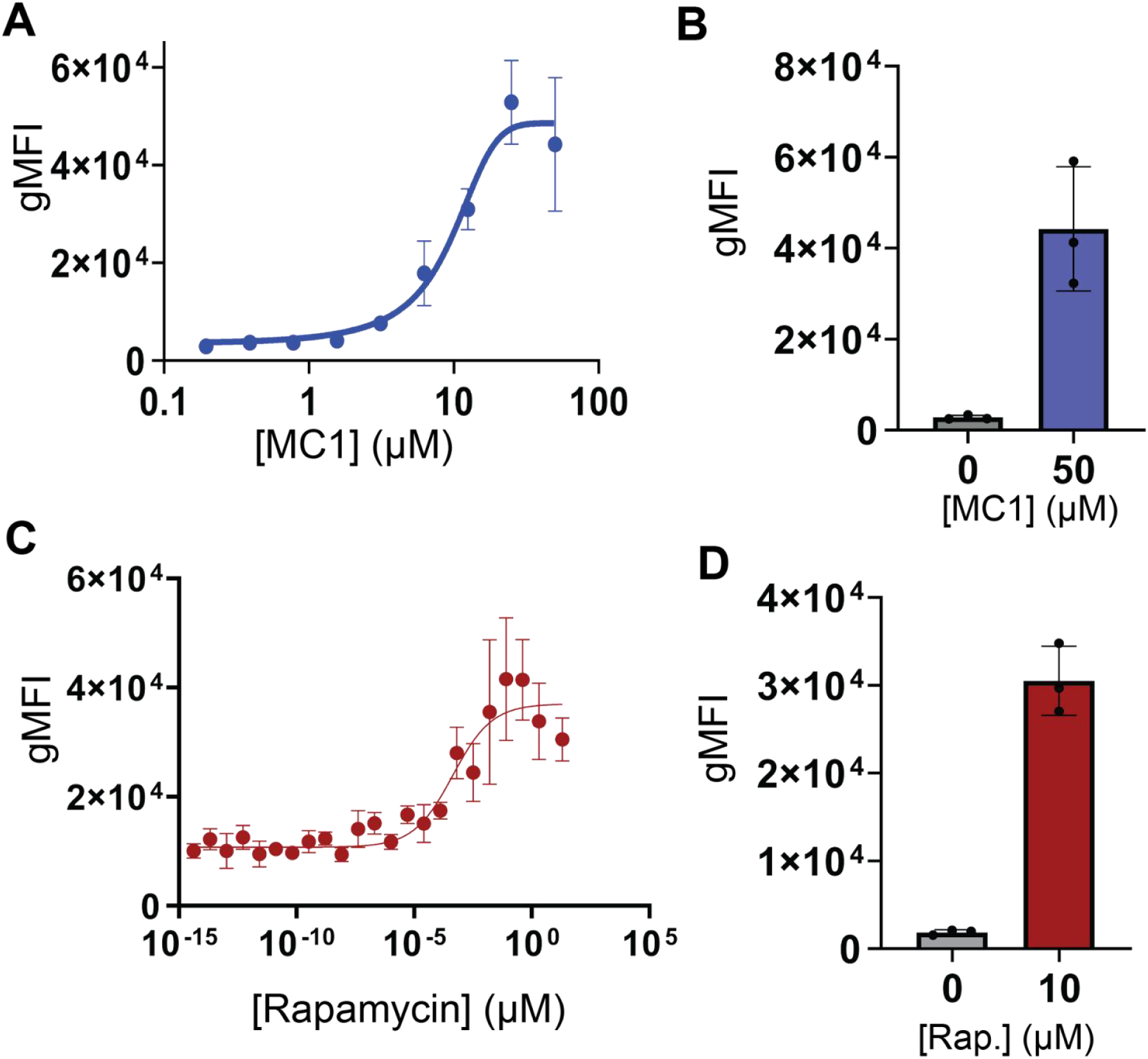
Designed CID system enables inducible transcription control. **a** HEK293T cells were transiently transfected with AD-CID7, DBD-CID7, and reporter. Cells were incubated with MC1 titrated in a 1 in 2, serial dilution with a starting concentration of 50 μM. Each concentration point plotted is the average of n=3 biological replicates from individual transfections. Error bars indicate the standard deviation from the mean of eGFP geometric mean fluorescence intensity (gMFI). **b** eGFP gMFI of cells transfected with DBD-CID7, AD-CID7, and reporter plasmids treated with either 0 μM MC1 (grey) or 50 μM (blue), Error bars represent standard deviation of the mean. n=3 biological replicates from individual transfections were plotted for both control and ligand treated samples. **c** HEK293T cells were transiently transfected with DBD-FRB, AD-FKBP, and reporter were incubated with rapamycin titrated in a 1 in 5, serial dilution with a starting concentration of 10 μM. Each concentration point plotted is the average eGFP gMFI of n=3 biological replicates from individual transfections. Error bars indicate the standard deviation from the mean. **d** eGFP gMFI of cells transfected with DBD-FRB, AD-FKBP, and reporter plasmids treated with 0 μM rapamycin (grey) or 10 μM (red), Error bars represent standard deviation of the mean. n=3 biological replicates from individual transfections were plotted for both control and ligand treated samples.

## Discussion

The CID system described enables inducible control of gene expression using a designed macrocyclic peptide with a designed protein pair. Very little experimental optimization was required to generate our CID system as the sequences of both the MC ligand and the designed protein it dimerizes are directly from Rosetta design calculations. Cell based assays did not show detectable toxicity of MC1, and the MC1 permeability in the PAMPA assay is in line with other membrane permeable compounds. The CID7-MC1 system signal, however, could be limited by the ability of the designed proteins to form homodimeric species with each construct and we observe some background dimerization both in the transcriptional assay as well as the Nanobit assay. Thus, design optimization to improve EC_50_ and reduce background homodimerization could further increase the utility of the system.

More generally, the ability to jointly *de novo* design both sides of a CID system opens the door to an almost unlimited number of new systems. These systems could act in concert for simultaneous yet independent control over many aspects of cell function, for both mechanistic investigation and external guidance of adoptive cell therapies.

## Methods

### Computational Design of macrocyclic peptides

Macrocycles (MCs) were designed using a 4-step process that involves (1) generation of hydrogen-bonding loops, (2) identification of loops that can close themselves into C2 symmetric MCs, (3) design of a sequence that encodes the C2-symmetric MC conformation (4) and chemical synthesis.

For hydrogen bond loop generation, polyglycine/sarcosine loops of the were constructed and the phi/psi/omega torsion angles were sampled from a Ramachandran map to generate candidate loop conformations. Loops were constructed as G-(NMe-G)_n_-G, with 3, 4, or 5 *N*-methylglycines spacing the terminal glycine residues. Conformers of each loop were only saved if the sampled phi/psi/omega torsions angles along the loop resulted in the terminal glycines adopting a hydrogen bond like those at hydrogen bonding paris within beta-sheets. This resulted in X, Y, and Z loops for n=3, 4, and 5 respectively.

For each of the identified loop conformations, the relative transformers of the NCaC coordinate frames between the N- and C-terminal glycines were calculated, hashed, and stored in hash tables. Closed conformations of C2-symmetric peptides were found by identifying individual loops that shared N-to-C and C-to-N binned transforms within the hashtables. Closed conformations of the full MCs were generated using a pyrosetta script by creating two copies of the identified loop (loop_i_ and loop_j_) and aligning the N terminal glycine of loop_i_ onto the C terminal glycine of loop_j_ and the C terminal glycine of loop_i_ onto the N terminalglycine of loop_j_. The resulting poses were merged to create a model of the MC. Each of the resulting MCs from this search contain two transannular hydrogen bonds.

Each identified backbone was subjected to a sequence design calculation using the FastDesignMover in Rosetta using a customized sequence composition that restricted the palette of amino acids to allow only hydrophobic residues. The resulting designed MCs were filtered to ensure the presence of the two transannular hydrogen bonds after the sequence design calculator. Surviving models were then clustered based on the ABEGO torsion bin and hydrogen bonding string^24^. A single low energy sequence per torsion hbond bin cluster was subjected to large scale energy landscape analysis using simple_cyc_pep_predict. Designs with funnel-shaped energy landscapes were prioritized for synthesis and structural studies.

### Protein design process

MC1 binding proteins were designed using a three step process that consisted of (1) RifGen (2) RifDock (3) Interface Design (Supplementary Note 1). Two design rounds were performed; the first with input scaffold sequences that were designed entirely with RosettaFastDesign^20^ and the second where the input scaffold sequences were redesigned with ProteinMPNN.

### Design of MC1 binding proteins using entirely Rosetta-derived scaffolds

Following the three-step design process, resulting designs were filtered for rosetta metrics (1) hydrogen bonds to the peptide backbone polar atoms, (2) interface shape complementarity, (3) few or 0 buried unsatisfied hydrogen bonds, and (3) ΔΔG. 19 designs were ordered for gene synthesis by Integrated DNA Technologies, the proteins expressed and purified from E. coli and binding assessed by equilibrium dialysis. We obtained a single binding hit, D_3_633_8x and then solved the co-crystal structure of D_3_633_8x in complex with MC1. The peptide was present in the designed binding pocket, but the atomic details were inaccurate as the peptide rotated such that it was bound in an asymmetric manner (Figure S2B).

### Design of MC1 binding proteins using MPNN redesigned scaffolds

In brief, ProteinMPNN was used to generate ten new sequences for each previously designed homodimeric scaffold backbone. After MPNN sequence design, structures were created with AF2 and filtered for a C a RMSD less than 2 **Å** to the input model, resulting in approximately thirteen thousand structures for binder design with MC1. Rotamer interaction fields were generated for MC1 and then used to dock the peptide into the scaffolds. Docks were filtered by rosetta metrics (interface area above 900, interface shape complementarity greater than 0.7, ΔΔG less than -40, contact molecular surface above 480, contact patch above 440). The protein peptide interface of docks passing these metrics was then redesigned with Rosetta FastDesign. The resulting designs were filtered for contact molecular surface above 460, ΔΔG less than - 40, at least three hydrogen bonds between the ligand and protein, interface shape complementarity above 0.72, interface area above 959, and for designs with less than one buried unsatisfied hydrogen bond. An AF2 structure prediction was generated and designs were filtered for pLDDT above 90, RMSD less than 1.1 **Å** between the design and prediction model, and a pTM score greater than 0.8, resulting in 263 designs.

### Protein expression and purification

Plasmids were constructed by either (1) ordering synthetic genes with an Avi tag from Integrated DNA Technologies (IDT) in pET-29b (+), resulting in the expression product: GLNDIFEAQKIEWHEGHHHHHHGSGSGENLYFQSGSGSSS[POI] or (2) ordering synthetic genes as a gblock from IDT then cloning the synthetic gene fragment into a custom entry vector with golden gate assembly (GGA). An Echo acoustic liquid handler (Beckman Coulter) was used to dispense 1 μL reaction volumes (0.024 μL water, 0.100 μL T4 ligase buffer (New England Biolabs), 0.600 μL eblock fragment at 4 ng/μl, 0.060 μL BSaI-HFv2 (New England Biolabs), 0.100 μL T4 ligase (New England Biolabs), 0.116 μL entry vector at 115 ng/μl). Reaction was placed in thermocycler (Biorad T100) with the lid heated to 105 °C and subjected to 37***°C*** for 20 min followed by 60***°C*** for 5 min. The custom vector, LM627 (addgene 191551), yields the expression product MSG[POI]GSGSHHWGSTHHHHHH^25^. The custom vector containing maltose binding protein (MBP) produces the expression product SHHHHHHG[MBP][POI]GS. 1 μL of plasmid was transformed in BL21 *E. coli* cells (NEB) and cells were incubated on ice for 20 min then heat shocked for 10 s. Cells were recovered in 100 μL SOC media for 1 h at 37***°C*** then plated onto lysogeny broth (LB, MpBio) agar plates under kanamycin antibiotic selection (50 µg/mL, Teknova) and grown overnight at 37***°C***. A single colony was inoculated into terrific broth (TB) II autoinduction media (MpBio) containing 50×5052 (glycerol 25% (v/w), glucose 2.5% (v/w), α-lactose 10% (v/w)), 2 mM MgSO_4_, and trace metal for overnight protein expression at 37***°C***. After approximately 20 h cells were harvested by centrifugation (4000 g, 5 min), resuspended in cell lysis buffer (20 mM Tris, 300mM NaCl, 25 mM imidazole, pH 8.0, 0.1 mg/mL lysozyme, 0.01 mg mL-1, Deoxyribonuclease I (DNAse I) and 1 mM Phenylmethylsulfonyl fluoride (PMSF), and sonicated using Qsonica, Q500 with a 4-pronged horn and 10 s ON, 10 s OFF, 45% amplitude for 5 min total ON time. Cellular debris was pelleted by centrifugation (16000 g for 25 min) and the proteins were purified from lysate supernatants by immobilized metal affinity chromatography (IMAC). Lysates were incubated with 500 μL of nickel-nitrolacetic acid (Ni-NTA) beads (Qiagen) in a 50 ml column (Biorad) and then washed with 3 column volumes (CV) of wash buffer (20 mM Tris, 300mM NaCl, 25 mM imidazole, pH 8.0). Proteins were eluted in 1.2 mL of elution buffer (20 mM Tris, 300mM NaCl, 300 mM imidazole, pH 8.0).

Proteins were then purified by SEC using an Akta Pure with an autosampler (Cytiva) on a Superdex S200 Increase 10/300 GL column (Cytiva 28990944) at room temperature at a flow rate of 0.75 ml/min. Proteins were purified into 25 mM Tris, 100 mM NaCl, pH 8.0 (SEC buffer). Fractions of interest were pooled, protein identity validated by LC-MS. Intact mass spectra was obtained via reverse-phase LC/MS on an Agilent G6230B TOF on an AdvanceBio RP-Desalting column, and subsequently deconvoluted by way of Bioconfirm 10.0 using a total entropy algorithm. Protein sequences, their observed and calculated masses are summarized in Supplementary Table 3.

For X-ray crystallography, the SNACtag was cleaved in solution^26^. Following IMAC purification, protein was exchanged into SNAC cleavage buffer (100 mM CHES, 100 mM NaCl, 100 mM acetone oxime, 500 mM guanidine HCl, pH 8.6). 2 mM NiCl2 was added and the protein solution was incubated overnight at 37°C. Solutions were then incubated with Ni-NTA resin to bind uncleaved products and the flow through was collected.

### MC synthesis

MCs were synthesized via solid phase peptide synthesis by WuXi Apptec using standard Fmoc-based solid-phase peptide synthesis on CTC resin with either HBTU/DIEA or HATU/DIEA coupling conditions. Linear peptides were cleaved from the resin and cyclized in solution followed by purification on reverse-phase HPLC^17^.

### MC1 characterization

Lyophilized peptide was dissolved in 50:50 ACN : water with 0.1% TFA. Sample was injected onto Agilent 1260 Infinity II HPLC equipped with a VWD detector and a Higgins Analytical PROTO 300 C18 5um 4.6×150 mm column (Agilent RS-1546-W185). Method used was 5-95%B over 45 minutes. Running buffers consisted of (A) water with 0.1% TFA and (B) 90% ACN with 0.1% TFA. Absorbance was collected at 214 nm. (Supplementary Fig. 10a).

Lyophilized peptide was dissolved in 50:50 acetonitrile (ACN) : water with 0.1% trifluoroacetic acid (TFA). Mass spectra was obtained via reverse-phase LC/MS on an Agilent G6230B TOF on an AdvanceBio RP-Desalting column with a gradient of 5-95% B over 2 minutes and flow rate of 0.5 ml/min. Running buffers consisted of (A) water with 0.1% TFA and (B) 90% ACN with 0.1% TFA. (Supplementary Fig. 10b).

### MC Crystallography

MC1 crystals were grown via vapor diffusion of Pentane into ethylacetate. Briefly, 1-3 mg of MC1 was dissolved in approximately 200 μL of ethyl acetate in a dram vial.

The dram vial was placed into a scintillation vial containing approximately 3 mL of pentanes. All other MC crystals were grown via vapor diffusion of water into acetonitrile (ACN). Briefly, 1-3 mg of macrocycle was dissolved in approximately 200 μL of ACN in a dram vial. The dram vial was placed into a scintillation vial containing approximately 3 mL of water. The scintillation vials were sealed and monitored for crystal growth. Crystals diffraction data were collected from a single crystal at synchrotron (on APS 24ID-C) and at 100 K. All crystal diffraction data was collected to 0.83 Å resolution. Unit cell refinement, and data reduction were performed using XDS and CCP4 suites^27, 28^. The structure was identified by direct methods and refined by full-matrix least-squares on F^2^ with anisotropic displacement parameters for the non-H atoms using SHELXL-2018/3^29,30^. Structure analysis was aided by using Coot/Shelxle^31,32^. The hydrogen atoms on heavy atoms were calculated in ideal positions with isotropic displacement parameters set to 1.2 × U_eq_ of the attached atoms. All designed MCs in this paper with solved crystal structures are summarized in Supplementary Table 1 and were deposited in Cambridge Crystallographic Data Centre (CCDC) with deposition numbers 2354604 (MC1, ethyl acetate conditions), 2354605 (MC1, ACN conditions), 2354606 (MC3), 2354607 (MC2), 2354608 (MC4), 2354609 (MC5), and 2360904 (MC6), and 2354610 (MC6-L4F).

### X-ray crystallography of CID7-MC1 complex

Purified protein CID7 with and without its ligand, MC1, was initially tested for crystallization via sparse matrix screens in 96-well sitting drops (200 nL drop volumes versus 95 μL reservoir volumes) using a mosquito crystallization robot (TTP LabTech). Crystallization conditions were then optimized with constructs that proved capable of crystallizing in larger 24-well hanging drops corresponding to initial mixtures of 1 μL well solution and 1 μL protein solution equilibrated against 1000 μL reservoirs. Crystals were grown in the presence of the ligands.

The crystal of the designed protein CID7 was grown from 100 mM sodium acetate pH 4.5 and 8% PEG 4000 at a protein dimer concentration of 153 μM. The ligand MC1 was added to the protein CID7 at a concentration twice the protein dimer and 3% final DMSO and incubated for 20 minutes before being added to crystal trays. The crystal was transferred to a solution containing 25% PEG 4000 and flash-frozen in liquid nitrogen. Data were collected at ALS Beamline 5.0.1 and processed using the program HKL2000^33^.

The structure of CID7 in complex with MC1 was solved by molecular replacement with Phaser (1.20) via PHENIX using the coordinates of the computationally designed structure as a search model^34, 35^. The structure was then built and refined using Coot (0.9.6) and PHENIX (1.20), respectively^37, 35^. The protein and solvent were modeled and refined first (while avoiding the modeling of any atoms within the binding site), and then the circular peptide ligand was built into the unambiguous density, with the last step being the addition of any missing waters. MC1 energies were calculated using eLBOW via PHENIX^35, 37^. Final Ramachandran statistics after refinement are listed in Supplementary Table 4. The co-crystal structure was deposited in the PDB with accession code 8VX7.

### X-ray crystallography of complex D_3_633_8x with MC1

Purified protein D_3_633_x8 with and without MC1 were initially tested for crystallization via sparse matrix screens in 96-well sitting drops (200 nL drop volumes versus 95 μL reservoir volumes) using a mosquito crystallization robot (TTP LabTech). Crystallization conditions were then optimized with constructs that proved capable of crystallizing in larger 24-well hanging drops corresponding to initial mixtures of 1 μL well solution and 1 μL protein solution equilibrated against 1000 μL reservoirs.

The crystal of the designed protein in the absence of ligand (“D_3_633_x8”) was grown from 2 M ammonium sulfate and 5% 2-propanol at a protein concentration of 216.92 μM. The crystal was transferred to a solution containing 2 M ammonium sulfate and 25% sucrose and flash-frozen in liquid nitrogen. Data were collected at ALS Beamline 5.0.2 at wavelength 400 nm and processed using program HKL2000^32^. The crystal was found to belong to a primitive orthorhombic space group (P212121) and yielded a 2 Å resolution data set.

The ligand ‘MC1’ was added to the protein construct D_3_633_x8 at twice the concentration of the protein dimer and 1.7% final DMSO and incubated for 5 minutes before adding to crystal trays. The crystal of the designed protein in the presence of ligand (“D_3_633_x8 MC1”) was grown from 2.5 M ammonium sulfate and 4% 2-propanol at a protein concentration of 43.38 μM and ligand concentration of 86.76 μM. The crystal was transferred to a solution containing 2.5 M ammonium sulfate and 25% sucrose and flash-frozen in liquid nitrogen. Data were collected at ALS Beamline 5.0.2 at wavelength 400 nm and processed using program HKL2000^32^. The crystal was found to belong to a primitive monoclinic space group (P21) and yielded a 2.05 Å resolution data set.

The structures of D_3_633_x8 and D_3_633_x8 MC1 were solved by molecular replacement with Phaser via PHENIX using the coordinates of the computationally designed structure as a search model^34, 35^. The structures were then built and refined using Coot and PHENIX, respectively^36, 34^. For the MC1 bound structure, the protein and visible surrounding ligands and solvent were modeled and refined (while avoiding the modeling of any atoms within the binding site, which displayed unambiguous density for the bound circular peptide ligand from the first rounds of modeling onwards). The final rounds of model-building were focused on fitting MC1 into the unbiased density that remained in the binding pocket. MC1 energies were calculated using eLBOW via PHENIX^11, 9^. Final Ramachandran statistics after refinement were as follows (given as % preferred, % allowed, % outliers, respectively): D_3_633_x8: 99.33, 0.67, 0.0; D_3_633_x8 MC1: 100.0, 0.0, 0.0 in Supplementary Table 4.

### Initial binding screen by equilibrium dialysis

6 μM MC1 stock solution was prepared in SEC buffer with 5% DMSO. Protein stock solutions were prepared in SEC buffer at either 20 or 100 μM. 50 μL protein solution was placed on one side of a Rapid Equilibrium Dialysis (RED) Device Inserts, 8K molecular weight cut off (MWCO) dialysis cassette (Thermo Fisher 89309) and dialyzed against 300 μL of MC1 stock solution. Dialysis cassettes were placed on 48 well plate and shaken (300 rpm, 30***°C***) for 5 h. 20 μL aliquots of each side of the cassette were diluted with 140 μL 90% acetonitrile (ACN), 0.1% formic acid (FA). Protein was then crashed out by centrifugation at 10,000 g for 10 min. Presence of MC1 on either side of the dialysis cassette for each protein sample was determined by injecting aliquots onto a Waters I-class ultra-performance liquid chromatography (UPLC) equipped with an AB Sciex 5600 quadrupole time of flight (QTOF) mass spectrometer. A calibration curve was generated for MC1 and was used, along with the area integration values of the extracted ion current (EIC) of MC1 mass, to determine the concentration of MC1 on either side of the cassette for each experimental sample. EIC peaks were integrated with Analyst® TF 1.6 and MultiQuant™ Software. The area ratio of MC1 on each side of the dialysis cassette for each respective sample was used to assess whether MC1 was binding each respective protein. CID7 showed area ratios greater than one in both the 20 and 100 μM protein sample experiments, suggesting CID7 bound MC1 (Supplementary Fig. 3).

### Characterization of binding affinity by ITC

SEC purified proteins were diluted to 50 μM in 20 mM Tris, 200 mM NaCl pH 8, 5% DMSO. Peptide solution was prepared at 250 μM in buffer matching that of the protein sample and was titrated into the protein sample. A control of 20 mM Tris, 200 mM NaCl pH 8, 5% DMSO was run for each experimental sample. ITC experiments were performed on an automated Microcal PEAQ-ITC. All titrations were performed with 19 total injections, where the first injection contained 0.4 μL of MC1 solution and the subsequent 18 injections contained 2 μL MC1 solution. The Microcal PEAQ-ITC Analysis Software was used to fit curves for dissociation constant determination.

### Mass Photometry

CID7-MBP protein concentration was determined by taking an A280 reading on nanodrop. Protein was diluted to 300 nM in SEC buffer then serially diluted 1:2 in SEC buffer. Samples equilibrated for approximately 1.5 h before recording measurements. 7.5 μl of sample was added to respective wells of a 24-well gasket atop a glass slide. 2 min videos were collected with regular field of view in AcquireMP. Individual particles were measured and processed into mass distributions with DiscoverMP. 40 nM Beta-amylase which contains monomer (56 kDa), dimer (112 kDa) and tetramer (224 kDa) in equilibrium was used to generate a mass calibration and convert contrast values into mass values. AcquireMP software was used to render all mass distribution plots. Three technical replicates were run for each sample. 0.4 nM homodimer *K*_D_ reported is the average calculated *K*_D_ of the three technical replicates with 9.4 nM total protein concentration.

The following equation was used to calculate fraction dimer (*F*_*D*_):

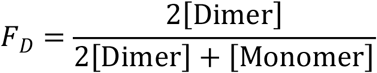

The following equations were used to estimate the *K*_D_ of the homodimer:

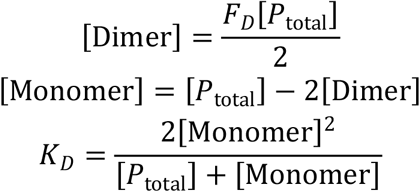

### Parallel Artificial Membrane Permeability Assay

PAMPA assay was conducted using a phospholipid pre-coated 96-well plate containing donor and acceptor compartments (Corning® BioCoat® Pre-coated PAMPA Plate System). Propranolol, a compound with high passive permeability^38^, was included as a positive control, whereas ranitidine, was used as low permeability control^39,40^.

Stock solutions of propranolol (20 mM), ranitidine (20 mM), and MC1 (20 mM) were prepared in PBS containing 5% DMSO. For the assay, 300 μL of each solution was added by triplicate to the designated wells of the donor plate. The acceptor plate was prepared by dispensing 200 μL of PBS with 5% DMSO into each well. The acceptor plate was then carefully positioned over the donor plate to allow the artificial membrane to contact the buffer solution. To minimize evaporation, the system was sealed with a lid and incubated for 17 h at room temperature on the benchtop.

Following incubation, 100 μL from each donor and acceptor well was transferred to two separate 96-well plates for analysis. Compound concentrations were determined by liquid chromatography–mass spectrometry (LC–MS) using a Xevo TQ Absolute XR Triple Quadrupole Mass Spectrometer (Waters Corporation) operated in multiple reaction monitoring (MRM) mode, which provides greater sensitivity by focusing on specific ion masses rather than performing a full scan.

Analyses were performed in positive ion mode using the following settings: capillary voltage, 3000 V; cone voltage, 50 V; desolvation temperature, 350 °C; desolvation gas flow, 800 L/h; cone gas flow, 150 L/h; and collision gas flow, 0.15 mL/min. Expected transition masses were monitored for analyte detection (Supplementary Table 2).

The mobile phases consisted of (A) 0.1% formic acid and 10 mM ammonium formate in MS-grade water, and (B) 0.1% formic acid in MS-grade acetonitrile. Chromatographic separation was achieved on an Acquity UPLC BEH C18 column (1.7 μm, 2.1 × 50 mm; Waters Corporation) with a flow rate of 0.3 mL/min and an injection volume of 2 μL. The gradient program was as follows: 10% B for 1 min, ramp to 100% B from 1 to 4.5 min, hold at 100% B for 1.5 min, then re-equilibrate to 10% B over 1.9 min prior to the next injection.

Calibration curves were generated from standard solutions at concentrations of 5, 7.5, 10, 15, and 20 mM (Fig. 2), and were used to quantify analyte concentrations in both donor and acceptor samples. Data processing and visualization of calibration curves were performed using the open-source software Skyline (25.1.0.237).

Equilibrium permeability coefficient (Pe) was calculated using the equations suggested by the manufacturer, based on the concentration difference between donor and acceptor wells over the incubation period, normalized to incubation time, membrane surface area, and sample volume.

Apparent permeability (Pe) was calculated using Corning’s mass-balance equation:

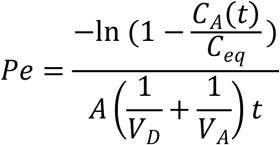

Equilibrium Concentration:

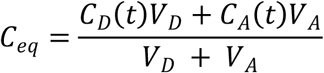

Mass Retention:

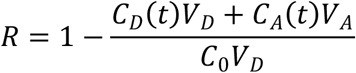

Where:

A = membrane filter

*V*_*D*_ = donor well volume

*V*_*A*_ = acceptor well volume

t = incubation time

*C*_*A*_*(t)* = analyte concentration in acceptor at time t (mM),

*C*_*D*_*(t)* = analyte concentration in donor at time t (mM),

*C*_*eq*_ = equilibrium concentration

*C*_*0*_= initial analyte concentration in donor well (mM)

For assays run with the donor solution placed in the receiver (bottom) plate (the PAMPA convention), Corning recommends a Pe cutoff of <15 nm/s (or 1.5 x 10^-6^ cm/s) for low permeability and >15 nm/s (or 1.5 x 10^6^ cm/s) for high permeability.

Data analysis for each analyte was conducted using Skyline, with calibration curves assessed using both linear and quadratic regression. For MC1, the quadratic model provided the best fit, reflecting the non-linear response of LC-MS detectors^41^.

### Cytotoxicity assay with MC1

100 μL of HEK293T (ATCC, CRL-3216) cells at a density of 0.2 mvc/cell were pleated onto a tissue culture treated 96-well plate and incubated at 37***°C*** until they reached 80% confluency. 10 mM MC1 in DMSO was serially diluted into cell media 1 in 4 from a starting concentration of 50 uM across 11 wells, resulting in a final concentration of 1% DMSO in all wells. Spent cell media was removed and replaced with MC1 serial dilution solution. Cell media was gently pipetted up and down before removing 2 uL aliquots from each well at 48 h ligand incubation timepoint. Sample aliquots were stored in 198 uL LDH storage buffer (200mM Tris-HCl (pH 7.3), 10% Glycerol (v/v), 1% BSA (w/v), sterile filtered) in a sterile 96 well tissue culture treated plate. The plates were sealed and stored at -20C prior to sample measurement on plate reader.

Cells were treated with cell media containing 1% DMSO for vehicle control. A maximum LDH control was run by treating cells containing cell media with 1% DMSO with 10% Triton X-100 (Millipore) 15 minutes prior to sample collection. Reagents from the LDH-Glo™ Cytotoxicity Assay kit (Promega J2380) were used for sample preparation. LDH standard samples were prepared following guidelines from the manufacturer and the standard curve generated was used to inform the sample dilution scheme.

Cell media in LDH storage buffer aliquots were thawed and 50 uL of each sample was transferred to a sterile white walled plate with clear bottom (Corning). 50 uL of LDH detection reagent obtained from the kit was added to each well and incubated with sample for 60 min prior to luminescence reading on a Synergy Neo2 plate reader with a LUM cube and with BioTek Gen5 software version 3.14. Percent cytotoxicity was calculated as suggested by the manufacturer using the formula below:

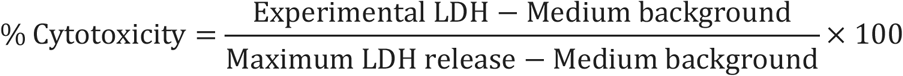

### Nanobit Assay in HEK293T cells

Plasmids were prepared by first digesting 1.5 µg of multiple cloning site vectors encoding either the larg ( ((e or small NanoBit protein (Promega N198A and N199A) with NheI-HF and EcoRI-HF (New England Bio Labs R3131 and R3101, respectively). CID7 genes were ordered as gblocks from Twist Biosciences with the manufacturer recommended base pairs added to ensure the fusion protein expressed has the adequate linker sequence (Supplementary Table 5). digested with NheI-HF and EcoRI-HF prior to ligation with the respective linearized plasmid. Respective linearized vector and digested CID7 genes were ligated with T4 DNA Ligase (New England Bio Labs) at room temperature for approximately 20 minutes and the enzyme was heat inactivated at 65°C for 5 min. Approximately 2 μL of each ligation reaction was added to 50 μL 5 a cells (New England Biosciences C2987), the cells rested on ice for 30 minutes, then were heat shocked at 42°C for 30 s. Cells were incubated with 200 μL SOC at 37°C for 1 h then plated on agar plates containing 50 μg/mL carbenicillin and allowed to incubate overnight at 37°C. Single colonies were grown in 5 mL LB with 50 μg/mL carbenicillin for approximately 20 h at 37°C. Colonies with the correctly assembled plasmid were purified by miniprep (Zymogen D4201) and sequence verified by whole plasmid sequencing (Plasmidsaurus).

HEK293T cells were plated onto 96-well TC-treated plates and allowed to reach 80% confluency prior to transfection. Plasmids stocks were diluted in Opti-MEM (Thermo Fisher 31985070) to a concentration of 6.25 ng/µl per construct. Viafect transfection reagent (Promega E4981) at 3:1 DNA : transfection reagent was added and the solution was mixed by pipetting up and down. Transfection solution sat for 10 min before it was added to each respective well.

Media containing transfection reagent was removed and 100 μL Opti-MEM with 2.5% DMSO was added to each well. Nano-Glo® Live Cell substrate was diluted 1 in 20 in Nano-Glo® LCS Dilution Buffer (Promega N2014), 25 μL was added to each well and luminescence was monitored for 20 min. 10 μL of either Opti-MEM with 2.5% DMSO (vehicle) or 2.4 uM MC1 in Opti-MEM with 2.5% DMSO was added to respective wells. Luminescence was measured continuously for another 1.5 h. To assess reversibility, cells were rinsed 3X with 100 μL vehicle, 110 μL vehicle was added to each well post rinsing, and then 25 μL of diluted substrate was added. Luminescence was recorded for another 30 min. All luminescence readings were recorded on a Synergy Neo2 plate reader with a LUM filter cube at 37°C. The maximum luminescence value occurring in each step of the experiment (initial addition of substrate, addition of compound or vehicle, compound washout and addition of more substrate) for each technical replicate was used to compute mean luminescence for plotting. All control samples received 10 μL vehicle at the compound addition stage. Fold difference in luminescence is calculated by dividing the mean luminescence signal of technical replicates after compound or vehicle addition by the mean luminescence signal of technical replicates before compound or vehicle addition.

### Transcriptional assay in HEK293T cells

Plasmids were prepared by ordering synthetic genes encoding the designed proteins from IDT as eBlocks, and contained overhangs for Golden Gate Assembly (GGA) into custom made entry vectors. These entry vectors were assembled by Gibson assembly from pre-existing plasmids. Constructed plasmid oligonucleotide sequences are enclosed in Supplementary Data File 1. They contained an ampicillin resistance gene as well as a ccdb lethal gene between the SapI cut sites. Subcloning into MA0005 results in a [POI]-zinc finger DNA binding domain expression product, while subcloning into MA0006 results in [POI]-p65 transactivation domain expression product. An Echo acoustic liquid handler (Beckman Coulter) was used to dispense 1 μL reaction volumes (0.130 μL water, 0.100 μL T4 ligase buffer, 0.425 μL eblock fragment at 4 ng/μl, 0.150 μL of SapI (New England Biolabs), 0.125 μL T4 ligase, 0.070 μL entry vector at 200 ng/μl). 1 μL of reaction mixtures were then transformed in 5-alpha competent *E. coli* cells (New England Biolabs) and cells were incubated on ice for 20 min then heat shocked for 10 s. Cells were recovered in 100 μL SOC media for 1 h at 37***°C*** and then plated on LB agar plates under carbenicillin selection with 100 µg/mL carbenicillin. A single colony of each design was inoculated into LB and the plasmids were grown overnight at 37***°C*** under carbenicillin antibiotic selection (100 µg/mL). Plasmids were then purified by miniprep.

An echo acoustic liquid handler was used to aliquot plasmid solutions into the respective wells of a non-tissue culture treated 384-well plate. Each well was filled to a final volume of 5 μL with nuclease free water and mixed with transfection reagent using a 24-channel pipette.

HEK293T cells were cultured in tissue culture-treated 384-well plates. Each well contained 50 µL of HEK293T cells at a density of 600,000 cells/mL in cell media (Dulbecco’s Modified Eagle Medium with Glutamax (DMEM, Thermo Fisher), 10% fetal bovine serum, heat inactivated (FBS, Thermo Fisher), 1X penicillin strep media (10,000 U/ml, Thermo Fisher)). At approximately 80% confluency cells were then transfected with the respective vectors. Cells expressing CID7 were transfected with an equal ratio of plasmid (150 ng total plasmid) using 150 mM NaCl and 3X polyethylenimine (PEI) solution. After approximately 10 minutes the plasmid transfection reagent was added to cultured cells. Cells were incubated with plasmid and transfection reagent for approximately 24 h at 37***°C***. After incubation with the transfection reagent, media was removed and replaced with new cell media. MC1 was serially diluted 1 in 2 from 50 µM across nine wells in cell media with 1% DMSO. Control wells were made by adding DMSO to cell media to a final concentration of 1%. Cells were incubated with new media for approximately 24 h at 37***°C*** before analysis by flow cytometry.

Cells expressing rapamycin binding proteins were transfected using 150 mM NaCl and 3X Viafect transfection reagent. After approximately 10 minutes, the plasmid Viafect solution was pipetted onto HEK293T cultured cells. Following 48 h incubation with transfection reagent, the media was removed from cells and new media was added. Rapamycin (Millipore Sigma) dissolved in DMSO prior to addition to cell media and then serially diluted 1 in 5 from 10 µM across twenty-three wells in cell media with 1% DMSO. Control wells were made by adding DMSO to cell media to a final concentration of 1%. Cells were incubated with new media for approximately 24 h at 37***°C*** before analysis by flow cytometry.

To prepare cells for analysis with flow cytometry, media was removed from the cells and the cells detached by incubating with 40 μL Trypsin-EDTA (0.25%, Thermo Fisher) for approximately 10 min. 50 μL of cell media was added to each well and cells were then resuspended by pipetting. The gating strategy for flow cytometry data is outlined in Supplementary Fig. 8.

### Flow cytometry

Experiments were analyzed on an Thermo Fisher Attune NxT flow cytometer. For analysis of eGFP expression, cells were gated on live singlets. To ensure only cells that were double transformed with both design-containing plasmids were included in downstream eGFP quantification, cells were then gated for the expression of mscarlett-1 and mtagBFP2. Finally, cells were gated for eGFP fluorescence. Geometric mean fluorescence intensity was computed in Flowjo and exported for processing with GraphPad Prism v11.0.0.

### Visualization and figures

All structural images for figures were generated with The PyMOL Molecular Graphics System, Version 2.5.8, Schrödinger, LLC. Data were analyzed, plotted, and processed with python (3.9.6), matplotlib (3.7.1), seaborn (0.12.2), and pandas (2.0.2).

## Data Availability

The protein crystallographic data generated in this study have been deposited in the PDB under accession codes 8VX7 (https://doi.org/10.2210/pdb8VX7/pdb), 8TM9 (https://doi.org/10.2210/pdb8TM9/pdb), and 8TLP (https://doi.org/10.2210/pdb8TLP/pdb). The macrocycle crystallographic data generated in this study have been deposited in the Crystallographic Data Centre (CCDC) with deposition numbers 2354604 [https://www.ccdc.cam.ac.uk/structures/Search?ccdc=2354604], 2354605 [https://www.ccdc.cam.ac.uk/structures/Search?ccdc=2354605], 2354606 [https://www.ccdc.cam.ac.uk/structures/Search?ccdc=2354606], 2354607 [https://www.ccdc.cam.ac.uk/structures/Search?ccdc=2354607], 2354608 (MC4, [https://www.ccdc.cam.ac.uk/structures/Search?ccdc=2354608], 2354609 [https://www.ccdc.cam.ac.uk/structures/Search?ccdc=2354609], 2360904 [https://www.ccdc.cam.ac.uk/structures/Search?ccdc=2360904], and 2354610 [https://www.ccdc.cam.ac.uk/structures/Search?ccdc=2354610].

## Code Availability

MC design code and protein design code used in this publication can be found at https://files.ipd.uw.edu/pub/macrocycle_cid/code.zip.

## Acknowledgements

This work has been supported by the Defense Threat Reduction Agency Grant HDTRA1-19-1-0003 (S.H. and M.Y.S.), Juvenile Diabetes Research Foundation International (JDRF) grant # 2-SRA-2018-605-Q-R (P.J.S.), Spark Therapeutics / Computational Design of a Half Size Functional ABCA4 (B.W.), The Open Philanthropy Project Improving Protein Design Fund (B.W., B.C.), National Institute of Health (NIH) R01 (M.A.K., B.L.S.), NIH R35 (M.A.K., B.L.S.), NIH S10 OD028581 (M.A.K., B.LS.), NSF DGE1762114 (M.A.K.), National Institutes of Health’s National Institute on Aging, cooperative agreement U19AG065156 (D.R.H.), The Audacious Project at the Institute for Protein Design (D.R.H, A.K.B., D.B.), Howard Hughes Medical Institute (HHMI) (A.K.B., D.R.H., D.B.), The Nordstrom Barrier Institute for Protein Design Directors Fund GF129460 (M.A.), NIHs National Institute of Allergy and Infectious Disease grant R0AI160052 (B.C.), Higgins Family (M.Y.S.), NSF Grant CHE-1629214 (A.K.B.), National Institutes of Health’s National Institute of Allergy and Infectious Disease, grant R0AI160052 (A.K.B.), DARPA program Harnessing Enzymatic Activity for Lifesaving Remedies (HEALR) under award HR0011-21-2-0012 (A.K.B.). The crystallographic data was collected at the Northeastern Collaborative Access Team beamlines, which are funded by the National Institute of General Medical Sciences from the National Institutes of Health (P30). This research used resources of the Advanced Photon Source, a U.S. Department of Energy (DOE) Office of Science User Facility operated for the DOE Office of Science by Argonne National Laboratory under Contract No. DE-AC02-06CH11357. This work was supported in part by the Advanced Research Projects Agency for Health APECx Program Award No. 1AY1AX000036 (S.C. and X.L.) Bill & Melinda Gates Foundation INV-043758 (S. C. and C.M.).

## Author Contributions

S.H. designed, screened, and experimentally characterized homodimeric binders to MC1 with support from B.W., M.A., and M.Y.S. P.J.S. developed the macrocycle design methodology and generated designed macrocycles for structure determination experiments. M.A.K., A.K.B., and P.J.S. collected and processed data for structure determination experiments. C. M. and X. L. collected and processed PAMPA data. S. C. collected and processed cytotoxicity assay data. D.R.H. designed homodimeric scaffold proteins. B.C. and D.R.H. developed initial design and docking methodology. S.H. and D.B. wrote the first draft of the manuscript. D.B. and B.L.S. offered supervision throughout the project. All authors read and contributed to the manuscript.

## Competing Interests Statement

The authors report no competing interests.

## Notes

### Competing Interest Statement

The authors have declared no competing interest.

