## Supplementary Information for "De novo design of a macrocycle induced dimerization system for cellular control"

| Peptide | Design Sequence | Experimental Sequence | C-alpha RMSD (Å) | All atom RMSD (Å) | Calculated mass (g/mol) | Deposition number |
|---|---|---|---|---|---|---|
| MC1 | apL*F*apL*F* | apL*F*apL*F* | 0.63 | 1.89 | 912.6 | 2354605, 2354604 |
| MC2 | apL*F*FPa*l* | apL*F*FPa*l* | 0.68 | 1.16 | 912.6 | 2354607 |
| MC3 | fpL*a*APl*p* | fpL*a*APl*f* | 0.78 | 1.89 | 912.6 | 2354606 |
| MC4 | MPa*l*iM*Pa*P | XPa*l*iX*Pa*P | 0.39 | 0.63 | 941.6 | 2354608 |
| MC5 | MPl*a*ipA*a*L* | XPl*a*ipA*a*L | 1.34 | | 929.6 | 2354609 |
| MC6 | VPa*L*aLpL*a* | VPa*L*aLpL*a* | 1.44 | | 915.6 | 2360904 |
| MC6-L4F | VPa*F*a*LpL*a* | VPa*F*aLpL*a* | 0.46 | | 949.6 | 2354610 |

**Supplementary Table 1. Summary of designed macrocycles with solved x-ray crystal structures.** Crystal structures for seven designed peptides were solved. Lower case in sequence represents D-amino acids. An asterisk following an amino acid residue represents N-methylation of the amide. X represents norleucine. C-alpha RMSD was measured between the design model and crystal structure. All atom RMSD was measured between all heavy atoms of the design models and crystal structures. Molecular weight was calculated for the experimental sequences that were synthesized via solid phase peptide synthesis and crystallized. Cambridge Crystallographic Data Centre (CCDC) numbers are listed for each structure. Deposition numbers for MC1 are for acetonitrile and ethyl acetate crystallization conditions respectively.

Supplementary Note 1:
While MC1, MC2, MC3, and MC4 showed close alignment to the design model, MC5 and MC6 crystal structures were slightly different from the design model. One loop of MC5 is in agreement between the design model and crystal structure, whereas the other loop is misfolded (Figure S1A). MC6 was initially designed with leucine residues at positions 4, 6, and 8, and the crystal structure of this compound was solved. Like MC5, the crystal structure showed that one loop of the MC matched the design model whereas the other loop was not aligned. A L4F mutant compound resulted in a forward funnel with a lower energy landscape than the original sequence.

Thus, the mutated sequence was crystallized, and the backbone aligns with the design model (Figure S1B-C).

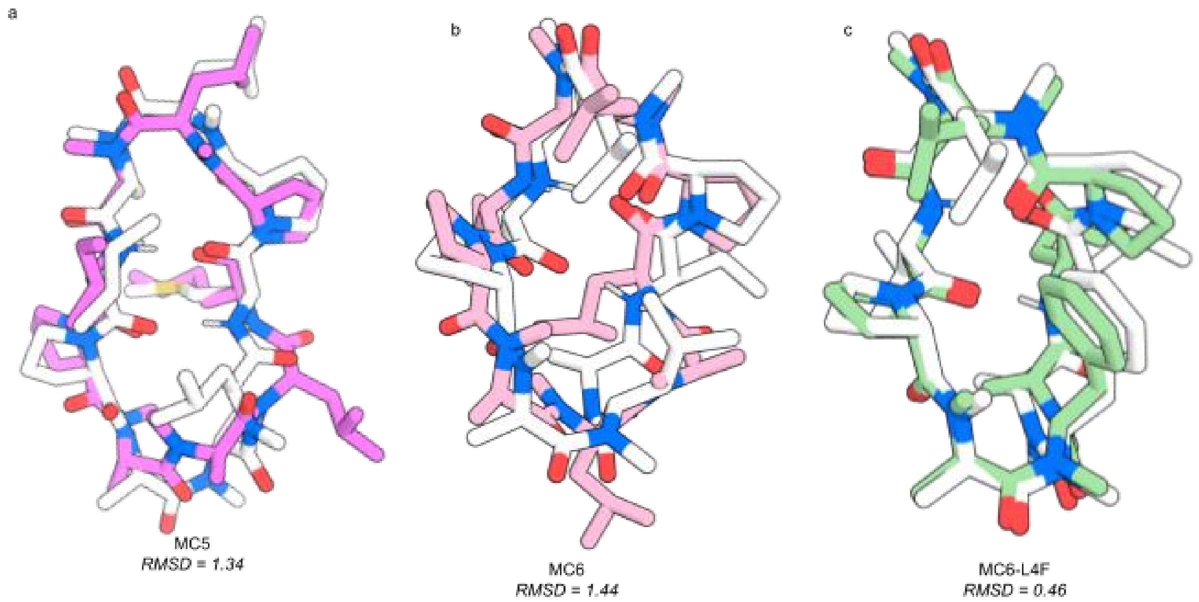

**Supplementary Figure 1. Design models and crystal structures of MC5, MC6, and MC6-L4F. a** Overlay of MC5 design model (white) with MC5 crystal structure (purple). **b** Overlay of MC6 design model (white) with MC6 crystal structure (pink). **c** Overlay of MC6-L4F design model (white) with MC6-L4F crystal structure (green). The C-alpha RMSDs between the design models and crystal structures are depicted beneath each MC.

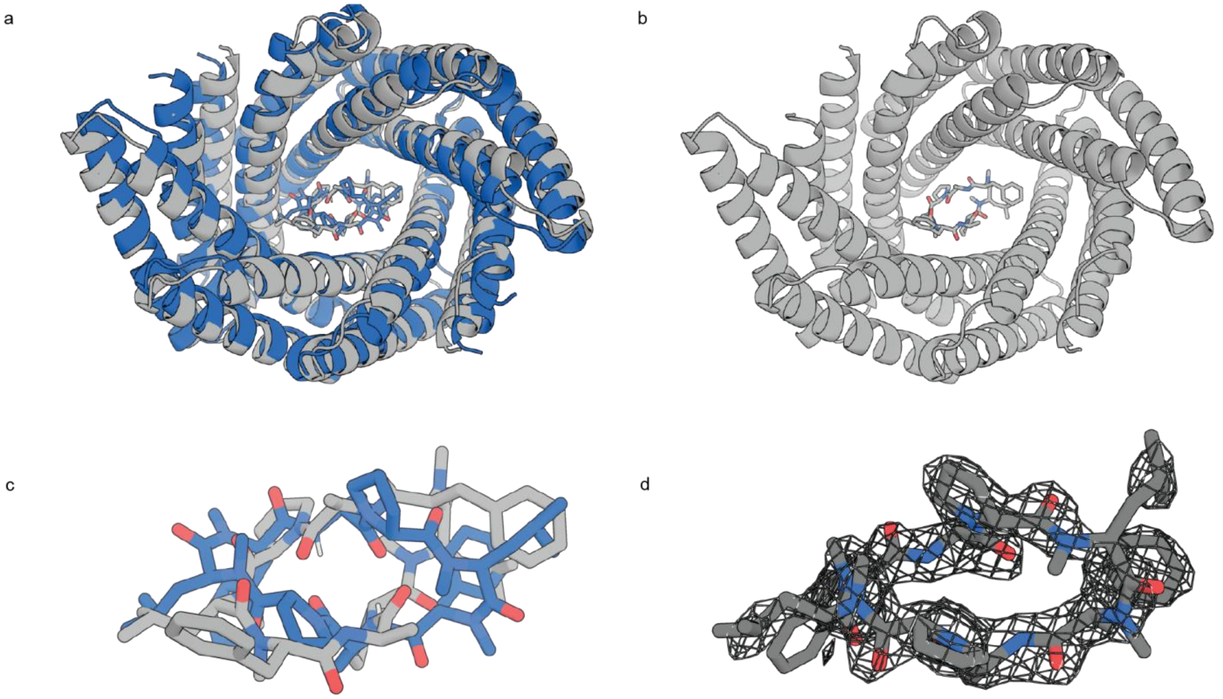

**Supplementary Figure 2. Overlay of the design model for original MC1 binding protein, D_3_633_8x, with the crystal structure. a.** Design model (blue) overlaid with the crystal strcture (grey) of the ligand bound protein. **b.** Crystal structure of the D_3_633 protein bound to MC1. **c.** Zoom in of MC1 of the design model (blue) overlaid with MC1 of the crystal structure (grey). **d.** Electron density map is shown in dark dray over MC1 of the crystal structure with contour level 1.

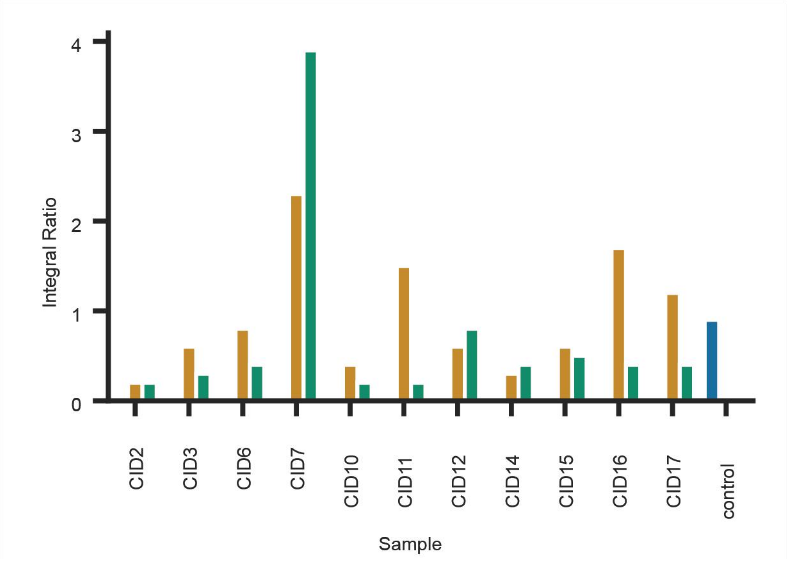

**Supplementary Figure 3. Initial assessment of MC1 binding by equilibrium dialysis.** Orange represents protein concentration of 20 µM, while green represents protein concentration of 100 µM dialyzed against 6 µM MC1. The control sample consisted of 25 mM Tris, 100 mM NaCl, pH 8.0 buffer with 0 µM protein dialyzed against 6 µM MC1. Integral ratio represents the ratio of MC1 concentration on the protein side of the cassette to MC1 concentration on the ligand side of the cassette. n=1 samples were used for all conditions tested.

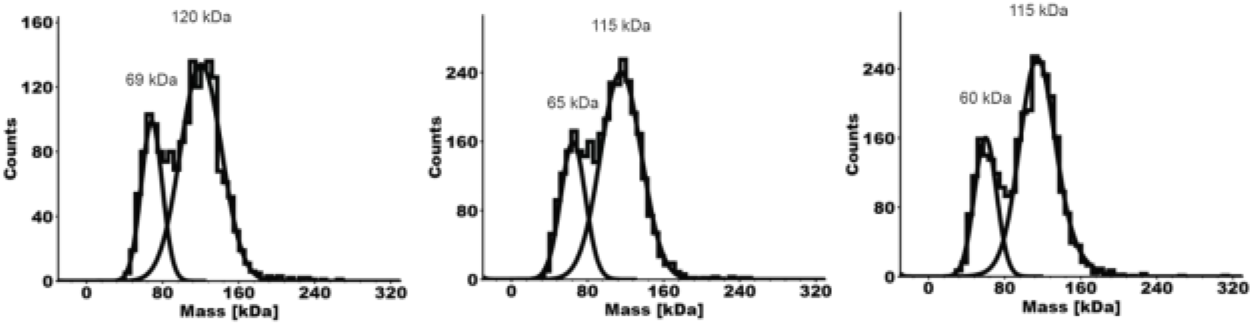

**Supplementary Figure 4. MBP-CID7 mass photometry distributions.** Mass distributions are shown for n =3 technical replicates containing 9.4 nM total protein. Masses obtained from converted contrast values are listed above each distribution.

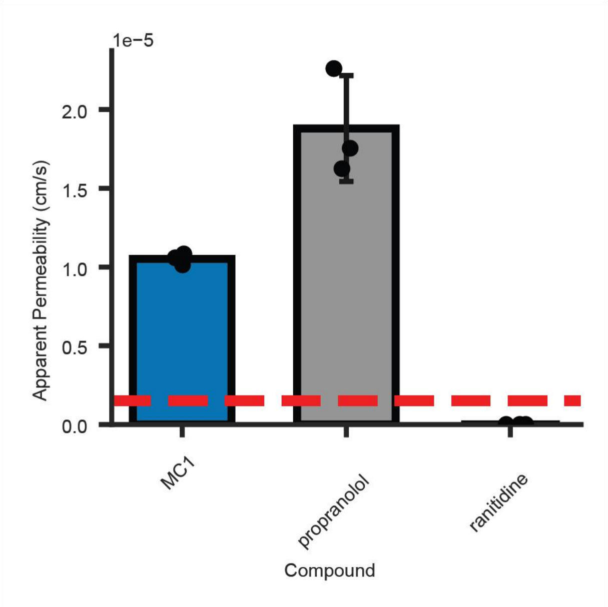

**Supplementary Figure 5. Membrane Permeability measurement of MC1.** Apparent permeability (Pe) was determined in PAMPA assay. Bar height represents mean Pe from n=3 technical replicates; error bars are the standard deviation of the mean calculated from the three technical replicates. Pe values were calculated from calibration curve data fit to quadratic regression. Ranitidine and propranolol were used as negative and positive controls for membrane permeability, respectively. The dashed, red line represents the Pe cutoff for a compound to be considered permeable. n=3 technical replicates were used for all conditions tested.

| **Analyte** | **Transitions** |
|---|---|
| Ranitidine | 315 >130 / 315 > 176 / 315 > 315 |
| Propranolol | 260 > 127 / 260 > 144 / 260 > 183 / 4: 260 > 260 |
| MC1 | 913.53 > 913.53 /930.53 > 930.53 ($NH_4^+$ adduct) / 935.51 > 935.51 ($Na^+$ adduct) |

**Supplementary Table 2. Calculated MS transitions for each analyte in the PAMPA expreiment.**

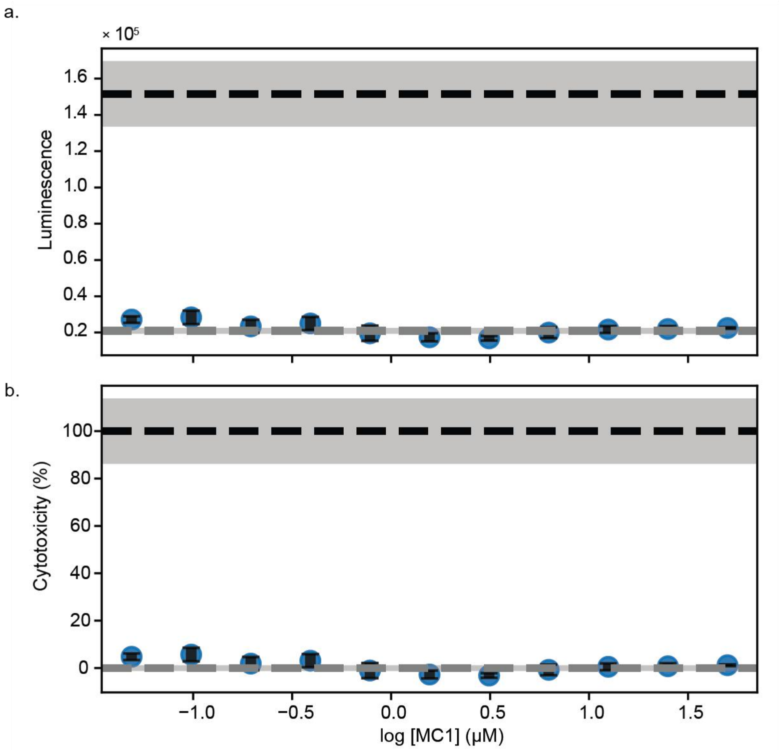

**Supplementary Figure 6. Percent cytotoxicity of HEK293T cells treated with MC1. a** Luminescence values were plotted against log MC1 concentration. The dashed horizontal line represents the mean of the max LDH release control samples consisting of 3 technical replicates. Data after 48 h incubation was MC1 is plotted. Error bars represent standard deviation of the mean. n=3 technical replicates were used for all conditions tested. **b** Percent cytotoxicity was calculated and plotted against log MC1 concentration. Percent cytotoxicity was calculated for each of three technical replicates after 48 h incubation with MC1. Error bars represent standard deviation of the mean. n=3 technical replicates were used for all conditions tested.

| **Protein** | **Expression Sequence** | **Observed mass (Da)** | **Calculated mass (Da)** |
|---|---|---|---|
| **CID7** | MSGMLEEIERLVLSGLLTGDKELLKKASELLKEEMEKLLEEGDLDALKKALQLAVNVADHNGDKELLAHAAEVIKRALDLALEAKDLQSAKYLASLALWIAKRAGDKELYAYLEEKIKKIIELAEEAGDRESLKILILLGIFIARDAGSEEVKAFVAEQLERLLGSGSHHWGSTHHHHHH | 20012 | 20012 |
| **MBP-CID7** | SHHHHHHGKIEEGKLVIWINGDKGYNGLAEVGKKFEKDTGIKVTVEHPDKLEEKFPQVAATGDGPDIIFWAHDRFGGYAQSGLLAEITPDKAFQDKLYPFTWDAVRYNGKLIAYPIA | 59567 | 59567 |

| | |
|---|---|
| | VEALSLIYNKDLLPNPPKTWEEIPALDKEL<br>KAKGKSALMFNLQEPYFTWPLIAADGGYA<br>FKYENGKYDIKDVGVDNAGAKAGLTFLV<br>DLIKNKHMNADTDYSIAEAAFNKGETAMT<br>INGPWAWSNIDTSKVNYGVTVLPTFKGQP<br>SKPFVGVLSAGINAASPNKELAKEFLENYL<br>LTDEGLEAVNKDKPLGAVALKSYEEELAK<br>DPRIAATMENAQKGEIMPNIPQMSAFWYA<br>VRTAVINAASGRQTVDEALKDAQTGSSGG<br>MLEEIERLVLSGLLTGDKELLKKASELLKE<br>EMEKLLEEGDLDALKKALQLAVNVADHN<br>GDKELLAHAAEVIKRALDLALEAKDLQSA<br>KYLASLALWIAKRAGDKELYAYLEEKIKKI<br>IELAEEAGDRESLKILILLGIFIARDAGSEEV<br>KAFVAEQLERLLGS |

**Supplementary Table 3. Summary and characterization of proteins expressed and purified from e. coli.** Observed mass represents the mass observed after deconvolution of the mass spectrum.

| | 8TLP | 8TM9 | 8VX7 |
|---|---|---|---|
| **Data collection** | | | |
| Space group | P $2_1$ $2_1$ $2_1$ | P 1 $2_1$ 1 | P $2_1$ $2_1$ $2_1$ |
| Cell dimensions | | | |
| *a*, *b*, *c* (Å) | 54.38, 55.03, 146.53 | 54.72, 54.56, 80.48 | 33.43, 79.80, 105.94 |
| ➪⭳ ⇦⭳ → (➡) | 90, 90, 90 | 90, 96.1, 90 | 90, 90, 90 |
| Resolution (Å) | 44.00 – 2.0 (2.071 – 2.0) | 47.4 – 2.05 (2.126 – 2.05) | 39.9 – 2.75 (2.848 – 2.75) |
| $R_{merge}$ | 0.056 (1.546) | 0.074 (0.42) | 0.113 (0.497) |
| *I* / ▽*I* | 31.36 (0.6) | 22.05 (2.07) | 23.44 (2.5) |
| Completeness (%) | 99.62 (96.41) | 99.18 (94.18) | 98.75 (91.03) |
| Redundancy | 11.8 (6.4) | 6.3 (4.8) | 11.6 (9.2) |
| **Refinement** | | | |
| Resolution (Å) | 2.0 | 2.05 | 2.75 |
| No. reflections | 30453 (2906) | 29557 (2789) | 7762 (700) |
| $R_{work}$ / $R_{free}$ | 23.3 / 28.1 (34.5 / 38.4) | 20.5 / 24.3 (25.2 / 29.2) | 25.5 / 30.4 (33.9 / 35.9) |
| No. atoms | | | |
| Protein | 450 | 452 | 322 |
| Ligand/ion | 5 | 93 | 66 |
| Water | 450 | 101 | 3 |
| *B*-factors | | | |
| Protein | 52.78 | 47.52 | 70.02 |

| | | | |
|---|---|---|---|
| Ligand/ion | 80.28 | 66.89 | 56.20 |
| Water | 53.06 | 46.83 | 52.18 |
| R.m.s. deviations | | | |
| Bond lengths (Å) | 0.007 | 0.002 | 0.001 |
| Bond angles (➡) | 0.81 | 0.40 | 0.36 |

**Supplementary Table 4. Final Ramachandran statistics after refinement for protein crystal structures deposited in the PDB.**

| Insert DNA sequence ordered | Entry Vector |
|---|---|
| GGACTGCTACGAATTCAATGCTGGAGGAGATCG<br>AGAGGCTGGTGCTTAGCGGCCTGCTGACAGGCG<br>ACAAGGAACTGCTCAAGAAAGCCAGCGAGCTG<br>CTGAAGGAGGAAATGGAGAAACTGCTGGAAGA<br>GGGCGACCTGGATGCCCTGAAGAAGGCTCTGCA<br>GTTGGCCGTGAACGTGGCTGACCACAACGGCGA<br>TAAGGAGCTGCTGGCTCACGCTGCCGAAGTGAT<br>TAAAAGAGCCCTGGACCTGGCCCTGGAGGCCA<br>AGGACCTTCAGAGCGCTAAGTACTTGGCCAGCC<br>TGGCTCTGTGGATTGCCAAGAGAGCCGGAGACA<br>AAGAGCTGTACGCCTACCTCGAGGAGAAGATC<br>AAGAAGATTATCGAGCTGGCCGAGGAGGCCGG<br>CGACAGGGAGTCTCTGAAGATCCTGATTCTGCT<br>GGGCATCTTCATCGCCAGAGACGCCGGAAGCG<br>AAGAGGTGAAGGCCTTCGTGGCCGAACAGTTG<br>GAGAGGCTGCTGTGAGCTAGCGGACGCTAC | pBiT1.1-N [TK/LgBiT] and pBiT2.1-N [TK/SmBiT] |

**Supplementary Table 5. DNA sequences ordered for split luciferase assay plasmid construction.**

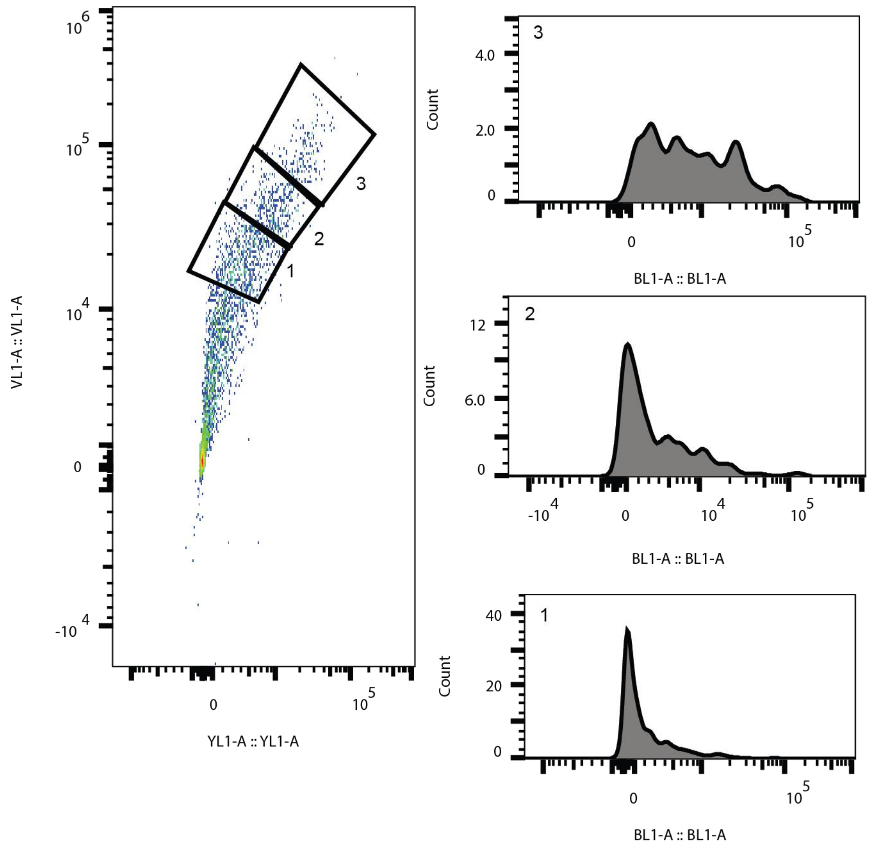

**Supplementary Figure 7. Construct expression vs GFP fluorescence of CID7. a** HEK293T cells were transfected with CID7-DBD, CID7-AD, and reporter plasmids and then treated with 1% DMSO. mscarlett-1 (assessed in YL1-A channel) vs mtagBFP2 (assesed in VL1-A channel) were plotted to show transfected population. Plots are composed of data from n=1 representative replicate. **b** eGFP fluorescence (assessed in channel BL1-A) distributions from gates 1, 2, and 3 in a.

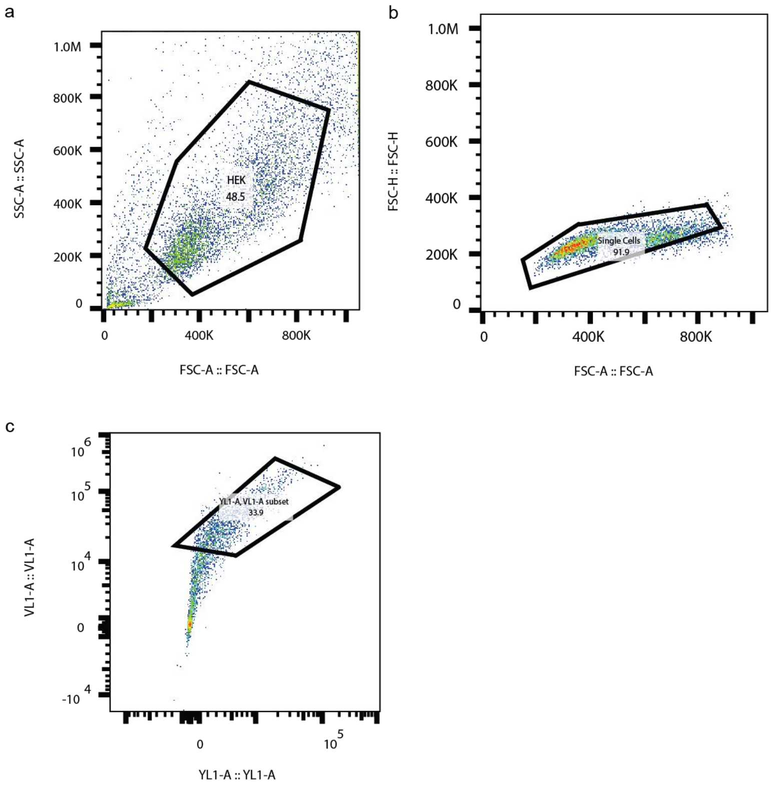

**Supplementary Figure 8. Gating strategy for HEK cells expressing construct proteins. a** Gates for live cells. **b** Live cells were then gated for singlet cells. **c** Of the singlet cells, the entire population expressing CID7 construct proteins as measured by RFP (channel YL1-A) and BFP (channel; VL1-A) expression markers. Plots are shown for one well of HEK cells on 384 tissue culture plate that were transfected with all three plasmids and treated with 1% DMSO in cell media. Gates were drawn in and plots were exported from flowjo software. Data represented is for cells expressing CID7 construct proteins.

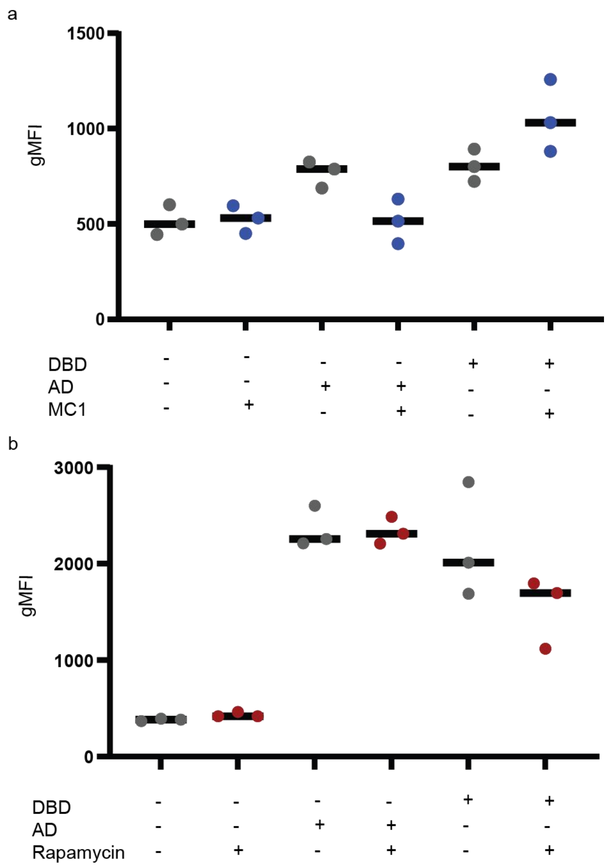

**Supplementary Figure 9. Construct controls for HEK cell experiments with either MC1 or rapamycin CID systems. a** HEK cells were transiently transfected with either 150 ng total plasmid of reporter and puc, DNA binding domain (DBD)-CID7, reporter, and puc, or activation domain (AD)-CID7, reporter and puc. Samples were treated with either 1% DMSO in cell media (grey) or 50 µM MC1 in cell media with 1% DMSO (blue). n=3 technical replicates were used for all conditions tested. is shown for each sample. **b** HEK cells were transiently transfected with either 150 ng total plasmid of reporter and puc, DBD-FRB, reporter, and puc, or AD-FKBP, reporter and puc. Samples were treated with either 1% DMSO in cell media (grey) or 1 µM MC1 (red). Data for three individual transfections is shown for each sample. Geometric mean of the GFP fluorescence for each sample was computed in flowjo and plotted in prism. n=3 technical replicates were used for all conditions tested.

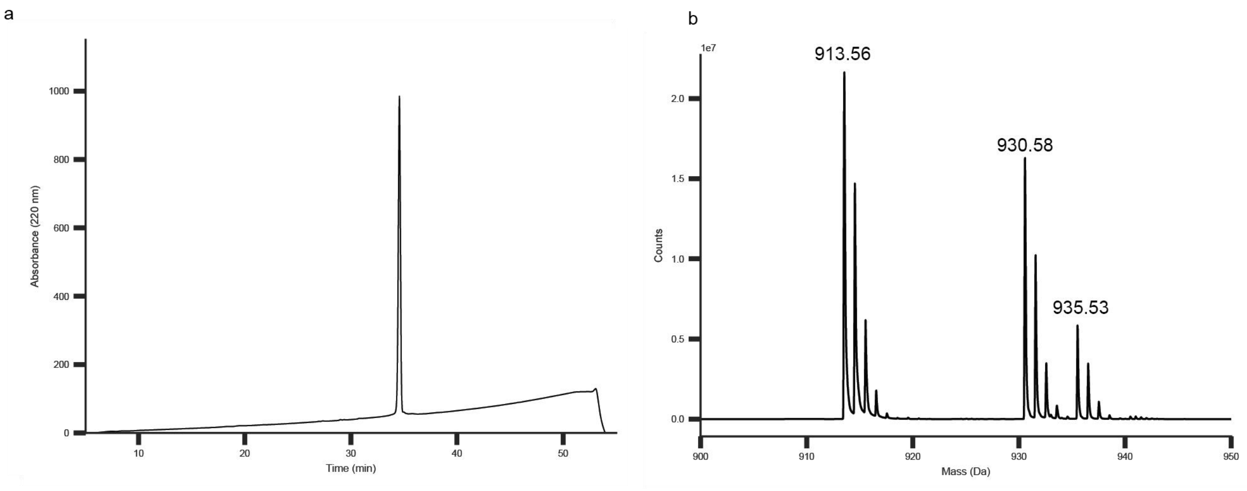

**Supplementary Figure 10. Purity trace and mass spectrum of MC1. a** Purity of the MC1 compound was 99% by HPLC. . **b** The observed mass from the mass spectrum corresponds to the calculated mass of 912.6 Da. 930.6 Da MS product corresponds to the loss of ammonia during ionization.

Supplementary Note 2

The following provides more background on the three step Rifgen, RifDock and Interface design process:

*RifGen*

RifGen was used to build disembodied hotspot side chains around the C2 symmetric peptide MC1, which had its symmetry axis aligned to the Z-axis and centered at the origin.

*RifDock*

We modified the inputs to RIFdock to only sample C2 docks. Then RifDock was used to dock the homodimeric scaffolds onto the MC1 hotspot interactions. To do this, we initially aligned the scaffolds symmetry axis to the Z-axis and centered them at the origin to match MC1. Instead of using RifDock's default hierarchical docking and scoring system that docks in 6D space, we provided C2 symmetric transforms to RifDock that moved the scaffolds up and down the Z-axis, while rotating them around the Z-axis, and performed a 180° flip perpendicular to the Z-axis to enumerate C2 symmetric docking orientations. These docks were scored with RifDock's default hash based scoring system and the top scoring docks were output for subsequent design.

*Interface Design*

The C2-symmetric MC1-scaffold complexes were then designed using the c2_peptide_design.xml script. Briefly, the script extracts an asymmetric unit from the input complex, applies rosetta C2 symmetry, add bonds to cyclize the peptide, designs the MC1-scaffold interface with FastDesign, performs FastRelax, and then repeats FastDesign, and FastRelax. The designed complexes were scored and various energy metrics were calculated by rosetta.
